# H2A.B is a cancer/testis factor involved in activation of ribosome biogenesis in Hodgkin Lymphoma

**DOI:** 10.1101/2021.01.19.427265

**Authors:** Xuanzhao Jiang, Jiayu Wen, Elizabeth Paver, Wu Yu-Huan, Gege Sun, Amanda Bullman, Jane E. Dahlstrom, David J. Tremethick, Tatiana A. Soboleva

## Abstract

Testis-specific regulators of chromatin function are commonly ectopically expressed in human cancers, but their roles are poorly understood. Examination of 81 primary Hodgkin Lymphoma (HL) samples showed that the ectopic expression of the eutherian testis-specific histone variant H2A.B is an inherent feature of HL. In experiments using two HL-derived cell lines derived from different subtypes of HL, H2A.B knockdown inhibited cell proliferation. H2A.B was enriched in both the nucleoli of these HL cell lines and primary HL samples. We found that H2A.B enhanced ribosomal DNA (rDNA) transcription, was enriched at the rDNA promoter and transcribed regions, and interacted with RNA Pol I. Depletion of H2A.B caused the loss of RNA Pol I from rDNA chromatin. Remarkably, H2A.B was also required for high levels of ribosomal protein gene expression being located at the transcriptional start site and within the gene body. H2A.B knockdown reduced gene body chromatin accessibility of active RNA Pol II genes concurrent with a decrease in transcription. Taken together, our data show that in HL H2A.B has acquired a new function, the ability to increase ribosome biogenesis.

## INTRODUCTION

Histones are essential nuclear proteins that compact eukaryotic DNA into chromatin, which provides a platform for the epigenetic control of genome function. Chromatin is built from nucleosomes, the universal repeating protein–DNA complex in all eukaryotic cells. A nucleosome comprises of two tight superhelical turns of DNA wrapped around a disk-shaped protein assembly of eight histone molecules (two molecules each of histones H2A, H2B, H3, and H4) (Luger et al., 2012). Different epigenetic mechanisms that alter the structure of the nucleosome to regulate genome function have been described. These include an extensive range of enzyme-catalyzed modifications of site-specific amino acid residues on the N-terminal tail of each histone, and altering the biochemical composition of a nucleosome by the substitution of one or more of the core histones with their variant forms (Buschbeck and Hake, 2017; Luger et al., 2012; Martire and Banaszynski, 2020). Collectively, this chromatin-based information at the genome-wide level is referred to as the epigenome.

Unlike core histones, which are transcribed only during S-phase, histone variants are expressed throughout the cell cycle (Kamakaka and Biggins, 2005). Among the core histones, the H2A family shows the greatest divergence in their primary sequence leading to the largest number of variants known. These variants include H2A.Z, H2A.X, MacroH2A, H2A.J, H2A.R and the “short histone” variants H2A.B, H2A.P, H2A.L and H2A.Q (Jiang et al., 2020; Malik and Henikoff, 2003; Molaro et al., 2018; Talbert and Henikoff, 2010). Key amino acid residue differences between the canonical histone H2A and its variant forms are strategically placed within the nucleosome and on its surface, and these differences affect nucleosome stability, higher-order chromatin compaction, and the interaction with reader proteins (Buschbeck and Hake, 2017; Doyen et al., 2006; Fan et al., 2004; Luger et al., 2012; Martire and Banaszynski, 2020; Shaytan et al., 2015; Zhou et al., 2007).

The short histone variants, designated as “short” because they lack an H2A C-terminus, are the most divergent. These histone variants appeared late in evolution in eutherian mammals, and are lineage-specific being expressed in the testis (Jiang et al., 2020; Molaro et al., 2018). The testis-specific expression of mouse H2A.B (H2A.B.3) is developmentally regulated with the peak of its expression occurring in highly transcriptionally active haploid round spermatids (the stage before transcription is switched off) (Anuar et al., 2019; Soboleva et al., 2012; Soboleva et al., 2014; Soboleva et al., 2017). By being targeted to the transcription start site (TSS) and intron-exon boundaries of active genes, H2A.B.3 is involved in enhancing transcription and the regulation of pre-mRNA splicing (Anuar et al., 2019; Soboleva et al., 2017). Notably, H2A.B.3 and H2A.L are RNA–binding proteins, which is consistent with the former being involved in splicing (Hoghoughi et al., 2020; Soboleva et al., 2017). *In vitro* chromatin reconstitution and transcription experiments have demonstrated that H2A.B (and H2A.B.3) decompact chromatin, thereby overcoming chromatin-mediated repression of transcription (Angelov et al., 2004; Soboleva et al., 2012; Zhou et al., 2007). Whether H2A.B can decompact chromatin *in vivo* is unknown.

Drastic alterations to the epigenome contribute to aberrant gene expression and have been shown to occur in virtually all cancer cell types (Rousseaux and Khochbin, 2009; Wang et al., 2011). However, the mechanisms responsible for these large-scale epigenetic abnormalities remain poorly understood. For about 30 years, it has been known that various somatic cancers display the aberrant activation of germ cell specific genes (known as cancer testis [C/T] genes), and many of these proteins are in fact regulators of the epigenome and chromatin structure (Debruyne et al., 2019; Simpson et al., 2005). It has been reported that H2A.B is abnormally expressed in Hodgkin Lymphoma (HL) cell lines but whether it is expressed in primary tumours has not yet been investigated (Sansoni et al., 2014; Winkler et al., 2012). In this study, we show for the first time that the expression of H2A.B is upregulated in all examined samples that represent all subtypes of primary HL (Figure 1).

**Figure 1.**
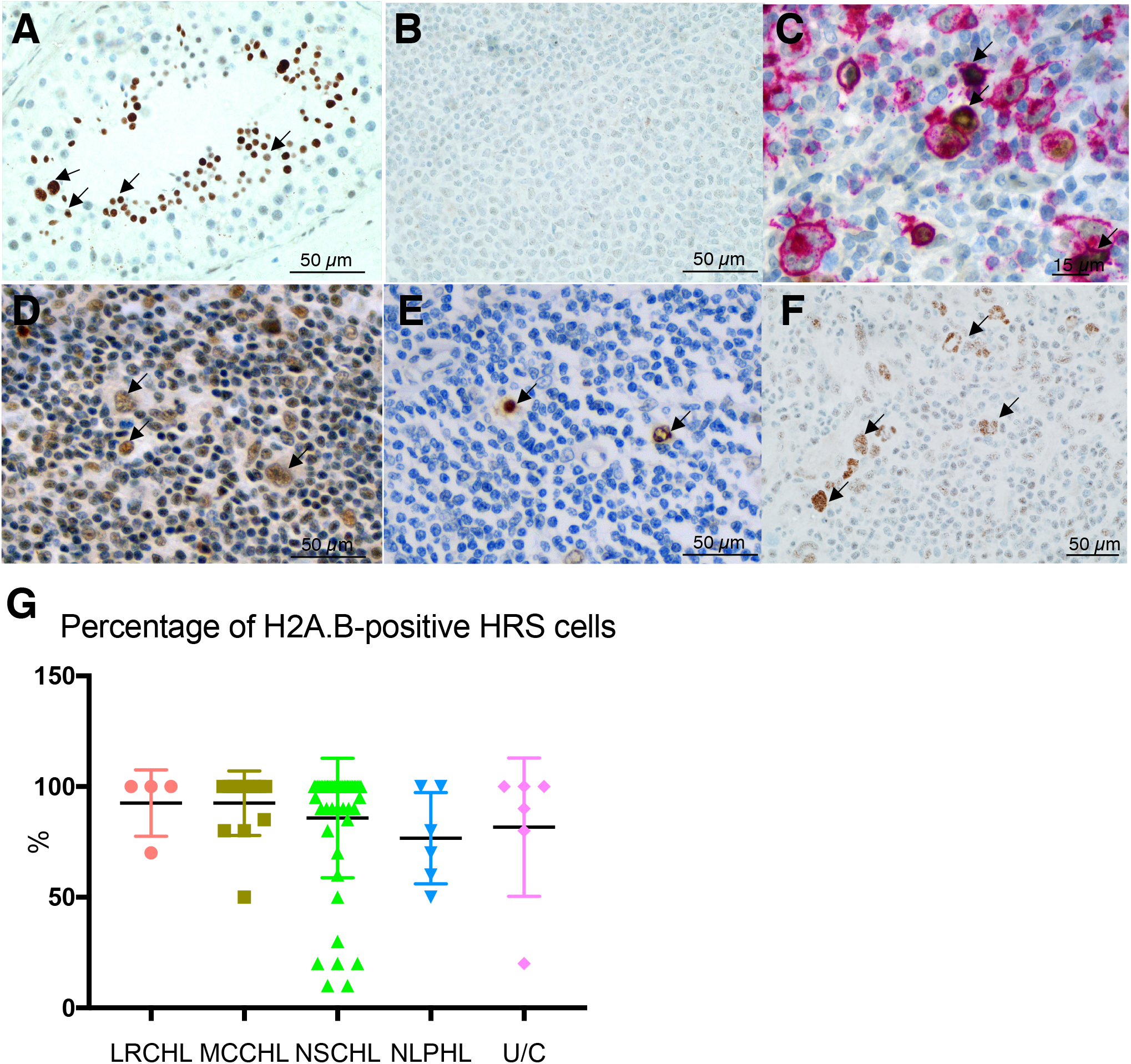
H2A.B is upregulated in all types of primary Hodgkin Lymphoma Immunostaining of primary HL with H2A.B (brown, indicated by black arrows) with a nuclear counterstain (blue). (**A**) Human testis (positive control). (**B**) Human tonsils (negative control). (**C**) Nodular sclerosis classical cHL, double staining with CD15 (pink) to identify HRS cells and H2A.B (brown). (**D**) Mixed cellularity cHL. (**E**) Lymphocyte rich cHL. (**F**) Nodular sclerosis HL. (**G**) Percentage of H2A.B-positive HRS cells in all 81 tumours. Lymphocyte Rich cHL (LRCHL), Mixed Cellularity cHL (MCCHL), Nodular sclerosis cHL (NSCHL), Nodular Lymphocyte Predominant HL (NLPHL), unclassified cases (U/C). Error bars are standard deviations, and the black line represents the average.

HL is a common hematopoietic malignancy that affects adults of all ages, and 10% of patients with HL do not respond to existing treatments (Shanbhag and Ambinder, 2018). The disease-causing cells of HL are mononucleated Hodgkin and multinucleated large Reed–Sternberg cells that arise from Hodgkin cells via endomitosis (Rengstl et al., 2013). Collectively, these cells are called Hodgkin Reed–Sternberg (HRS) cells. HRS cells are thought to arise from germinal centre B cells that differentiate into plasma cells through undergoing somatic hypermutation of the variable (V) regions of immunoglobulin genes upon antigen activation. Most mutations are disadvantageous and these crippled B cells undergo apoptosis. Sometimes, however, crippled B cells escape apoptosis and become precursors to HRS cells (Kanzler et al., 1996a; Kanzler et al., 1996b). Although HRS are clonal tumour cells, they are rare and represent only 1–2% of cells in tumour tissue. The rest of the tumour comprises of a mixed infiltrate containing non-neoplastic T cells, B cells, eosinophils, neutrophils, macrophages, and plasma cells (Pileri et al., 2002).

Having confirmed that H2A.B is expressed in the nuclei of cancerous HRS cells in primary HL samples (Figure 1); we hypothesized that H2A.B regulates the expression of genes that are required to promote oncogenesis and HL survival. To test this hypothesis, we established a Tet-inducible lentiviral short hairpin RNA (shRNA)–mediated H2A.B knockdown system in two different HL cell lines. Our experiments uncovered an unexpected function for H2A.B that is not observed in the mouse testis. Specifically, we show here that H2A.B displays a novel feature in enhancing ribosome biogenesis by increasing Pol I transcription of rDNA and Pol II transcription of ribosomal protein genes. These findings provide new insights into a well-established phenomenon that links hyperactive ribosome biogenesis with high cell proliferation rates and cancers.

## RESULTS

### H2A.B is ectopically expressed in primary HL tumours

To investigate whether upregulation of H2A.B expression is a feature of HL, we raised a rabbit polyclonal H2A.B-specific antibody and confirmed its specificity (Supplementary Figure 1) using the following approaches: 1) a Western a blot analysis; 2) immunoprecipitation of endogenous H2A.B; 3) immunofluorescence detection of H2A.B exogenously expressed in HEK293T cells; and 4) the observation that endogenous non-denatured H2A.B protein can be detected in paraffin-embedded human testis samples. As expected, pachytene spermatocytes and spermatids were positively stained (Soboleva et al., 2017) (Figure 1A, positive control). By contrast, H2A.B is not expressed in lymphocytes of normal tonsils and no signal was observed in these cells (Figure 1B, negative control).

We then examined 81 paraffin-embedded HL primary tumour samples that represented four subtypes of classical HL: 51 Nodular Sclerosis classical HL (cHL) samples, 14 Mixed Cellularity cHLsamples, 4 Lymphocyte Rich HL samples, 6 Nodular Lymphocyte Predominant non-classical HL samples and 6 cHL samples that were unclassified. The immunostaining results were unequivocal and showed H2A.B expression in all 81 samples and all HL subtypes (Figure 1C–G). Importantly, the staining was confined to the nuclei of cancerous HRS cells (Figure 1C, staining with anti-CD15 antibody identifies HRS cells, pink). On average, 87% of HRS cell nuclei within HL samples were positive for H2A.B (Figure 1G). Interestingly, unlike normal lymphocytes of disease-free lymphoid tissue (Figure 1B), infiltrating lymphocytes in HL samples displayed weak positive staining for H2A.B in many tumours (compare Figure 1B with Figures 1D and F). This observation raises the interesting possibility that infiltrating lymphocytes may also undergo transformation and H2A.B upregulation in the vicinity of HRS cells. Importantly, the presence of H2A.B in the nuclei was confirmed for all subtypes of HL (Figure 1G). In conclusion, we show that the ectopic upregulation of H2A.B gene expression in HL is not a sporadic event but an unequivocal nuclear characteristic of this tumour type. Given that primary HL tumours contain only 1–2% of HRS cells, we next used established HL cell lines derived from HRS cells to investigate whether H2A.B plays a role in HL carcinogenesis (Kapp et al., 1995).

### H2A.B is expressed and incorporated into chromatin in HL cell lines

We interrogated three HL cell lines L1236 (mixed cellularity, stage IV, B-cell lineage), L428 (nodular sclerosis, stage IVB, B-cell lineage), and HDML-2 (nodular sclerosis, stage IV, T-cell lineage) in order to confirm their identity, determine whether H2A.B carries any mutations, and investigate the level of its expression. We confirmed the cellular identity of these cell lines by examining their cellular morphology (data not shown) and by the reported expression pattern of four homeobox genes (Nagel et al., 2007); *PAX5* and *HLXB9* (expressed in all three cell types), and *PRAC1* and *PRAC2* (expressed only in L428 cells) (Supplementary Figure 2A).

The genomic DNA of the H2A.B-encoding genes was sequenced for all three HL cell lines, and the alignment with the *H2AFB1* gene showed no DNA mutations either in the coding region or in the 5’- and 3’-untranslated regions (Supplementary Figure 2B). Real-time qPCR confirmed the expression of H2A.B in all three cell lines and that the highest level of expression occurred in L1236 and L428 cell lines (Figure 2A) as reported previously (Sansoni et al., 2014). By contrast, the expression of H2A.B was low in HDML-2 cells, and it was not expressed in germinal centre B lymphocytes or in the Non-Hodgkin Lymphoma cell line, Ramos (Figure 2A). Western analysis confirmed the expression of H2A.B in L1236 and L428 cells (Figure 2B). The subcellular salt fractionation of L1236 cells into a tightly chromatin-bound fraction, a loosely chromatin-bound fraction, and a cytoplasmic–nucleoplasmic fraction demonstrates that H2A.B was incorporated into chromatin. A significant proportion of H2A.B is also not bound to chromatin, reflecting its highly dynamic chromatin binding properties (Bonisch et al., 2012; Gautier et al., 2004).

**Figure 2.**
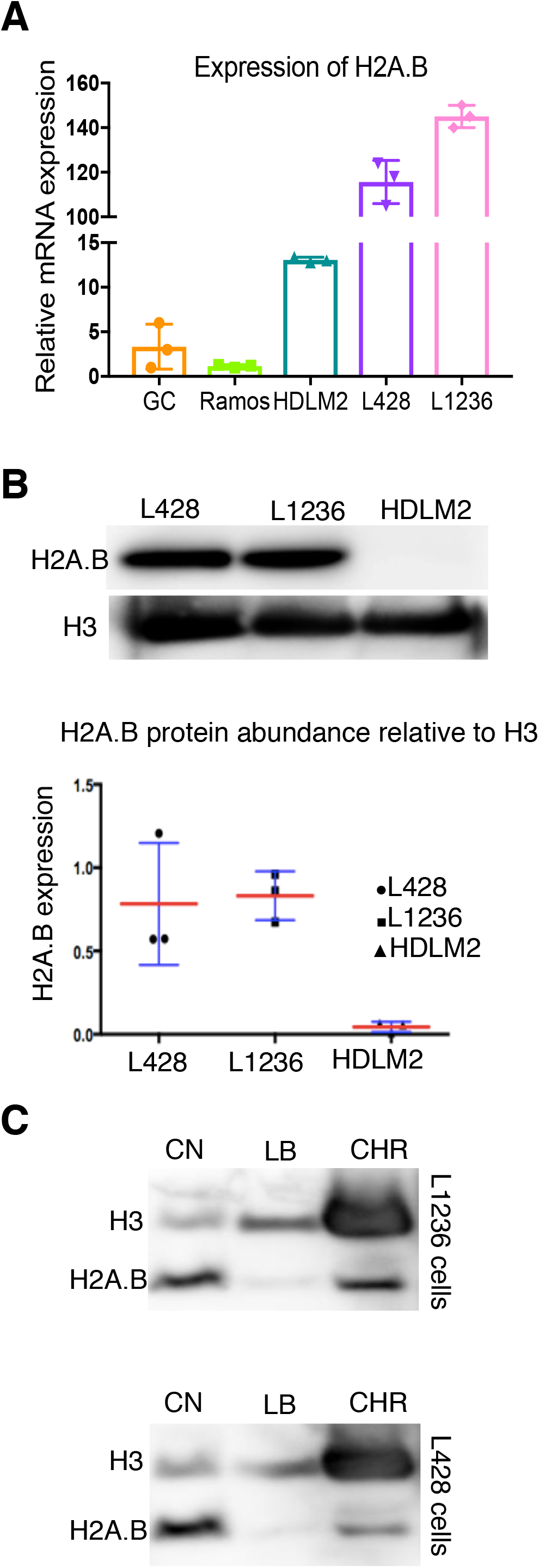
H2A.B is upregulated in Hodgkin Lymphoma cell lines. (**A**) Relative mRNA expression levels of H2A.B in L1236, L428, HDLM2 HL cell lines and Germinal centre B lymphocytes (GC) compared to the Ramos B lymphocyte cell line. (**B**) Western blot analysis of H2A.B protein expression relative to histone H3 protein expression. Error bars represent the standard deviation, and the red line shows the average. (**C**) Subcellular fractionation of L1236 and L428 cells using KCl salt. CN, cytoplasmic and nucleoplasmic fraction, LB, Loosely bound to chromatin, CHR, chromatin fraction. Note that the H3 Western blot signal in CHR is overexposed.

### H2A.B is enriched in rDNA chromatin and interacts with RNA Pol I

Unexpectedly, the H2A.B immunostaining pattern of L1236 cells revealed that this histone variant was enriched in the nucleolus, which was confirmed by the coimmunostaining for the nucleolar marker Fibrillarin (Figure 3A). Significantly, H2A.B was also found to be located in the nucleoli of HRS cells from a primary HL patient sample (Figure 3B). Given that H2A.B is not found in the nucleolus in mouse germ cells, this shows that this localization pattern is a tumour-specific phenomenon (Soboleva et al., 2017).

**Figure 3.**
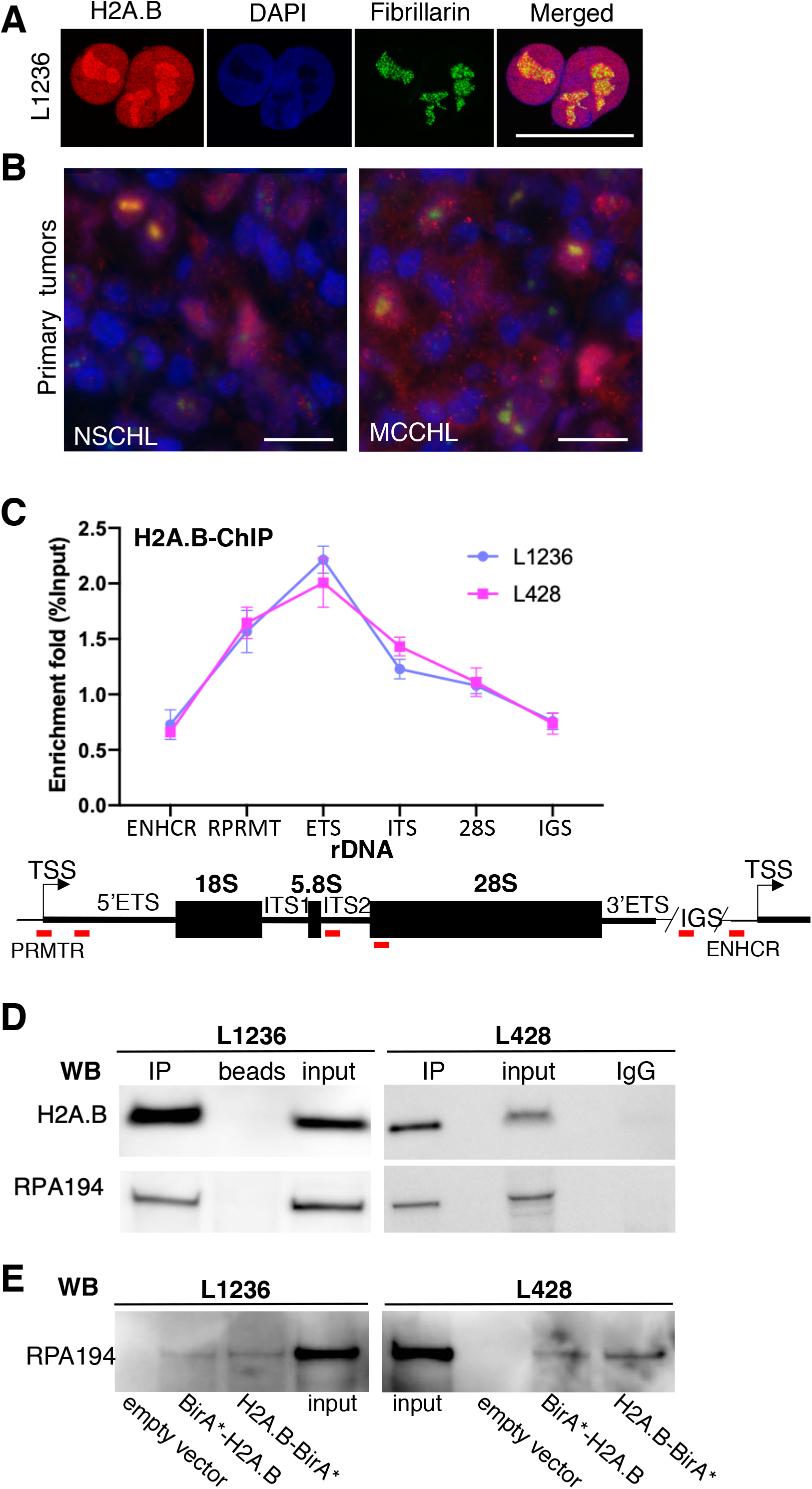
H2A.B is located at rDNA repeats. (**A**) H2A.B (red) is co-localized with fibrillarin (a nucleolar marker, green) in the nucleoli of L1236 cells. DAPI (blue) was used to counterstain nuclei. Scale bar is 10 μm. (**B**) Primary tumours stained with anti-H2A.B (red) and anti-fibrillarin (green) antibodies. Co-localization appears as a yellow signal, and DAPI (blue) counterstains nuclei. NSCHL, nodular sclerosis classical cHL; MCCHL, mixed cellularity cHL, scale bar is 25μm. (**C**) ChIP-qPCR showing the enrichment of H2A.B-containing nucleosomes at rDNA chromatin. ChIP with anti-H2A.B antibody was performed in L1236 and L428 cells followed by qPCR with rDNA primers corresponding to untranslated and spliced-out regions. Intergenic and enhancer regions were used as negative controls. Below the graph is a schematic diagram of an rDNA repeat. PRMTR, promoter; ETS and ITS, external and internal transcribed regions, respectively; IGS, intergenic spacer; ENHCR, enhancer region. Enrichment was calculated relative to input DNA. Red lines show the location of the amplification regions (PRMTR, 5’ ETS, ITS2, 28S, IGS and ENHCR). (**D**) H2A.B interacts with Pol I. Western blot showing co-immunoprecipitation of RPA194 with H2A.B. The Pol I subunit RPA194 was immunoprecipitated from L1236 (left panel) and L428 (right panel) nuclear extracts. The pull-down by RPA194 antibody (IP), pulldown by empty beads or IgG isotype control, and 1% input protein were loaded for Western blot detection using anti-H2A.B and anti-RPA194 antibodies. (**E**) H2A.B interacts with RPA194 in live cells. Western blot analysis was performed to detect RPA194 in the BioID samples from L1236 cells (left panel), and L428 cells (right panel). The samples were prepared using pull-down with streptavidin beads from the cell lysate of the empty vector, BirA*-H2A.B and H2A.B-BirA* transduced cells, which were preincubated with 50 μM biotin for 24 h. The positive control was loaded as 1% input.

This observation raised the intriguing possibility that H2A.B may have a role in rDNA transcription. To investigate this possibility, we first determined whether H2A.B-containing nucleosomes are found on rDNA by performing H2A.B ChIP-qPCR experiments. The qPCR primers were designed to span various regions of the rDNA repeat: the promoter (PRMTR), the 5’ external transcribed spacer (5’ETS), the internal transcribed spacer (ITS), the 28S subunit, the enhancer (ENHCR) as well as the intergenic spacer (IGS) as a negative control. Indeed, H2A.B-containing nucleosomes were found on rDNA being particular enriched at the promoter, 5’ ETS, and the ITS in both L1236 and L428 HL cell lines (Figure 3C).

Next, whether H2A.B interacts with RNA Pol I was investigated in order to provide evidence that H2A.B is located at actively transcribed rDNA repeats. First, immunoprecipitation of L1236 and L428 HL nuclear extracts with anti-RPA194 (a subunit of Pol I) antibodies followed by a H2A.B Western analysis showed that H2A.B co-immunoprecipitated with Pol I (Figure 3D). Second, to confirm that this interaction occurs in live cells, we employed a biotin labelling assay (BioID). In this assay, the biotin ligase BirA (which can label proteins in the vicinity of 10nm) was fused to either the N- or C-terminus of H2A.B and these fusion proteins were expressed in L1236 and L428 HL cells. H2A.B-BirA localised to the nucleolus (Supplementary Figure 3) where it was able to biotinylate RPA194 (Figure 3E). Therefore, H2A.B is in close proximity to Pol I implying that it does have a role in rDNA transcription.

### H2A.B increases rDNA transcription and regulates cell proliferation

To examine directly the involvement of H2A.B in rDNA transcription, we established a doxycycline inducible knockdown shH2A.B system in both L1236 and L428 HL cell lines, and tested two shH2A.B constructs were tested (shH2A.B and shH2A.B-2). Knockdown of H2A.B mRNA and protein expression with the shH2A.B construct was more efficient in L1236 (~95%) than in L428 (~75%) cells compared with the nontargeting shRNA (shNTC) control after 5 days of dox treatment (Supplementary Figure 4A–C).

Next, we compared the rRNA transcription levels in shH2A.B versus shNTC treated cells. Since the processed mature rRNA transcripts have a long half-life, the levels of primary transcripts (pre-rRNA) were determined by using qPCR primers that were designed to target the 5’ ETS and ITS. Unequivocally, the level of pre-rRNA transcripts was significantly reduced in both cell lines when the expression of H2A.B was inhibited (Figure 4A). Further, Pol I (RPA194) ChIP-qPCR experiments showed a significant reduction in Pol I at the 5’ ETS and ITS in H2A.B knockdown L1236 cells (Supplementary Figure 5). In can therefore be concluded that H2A.B is required for high levels of transcription in HL cell lines.

**Figure 4.**
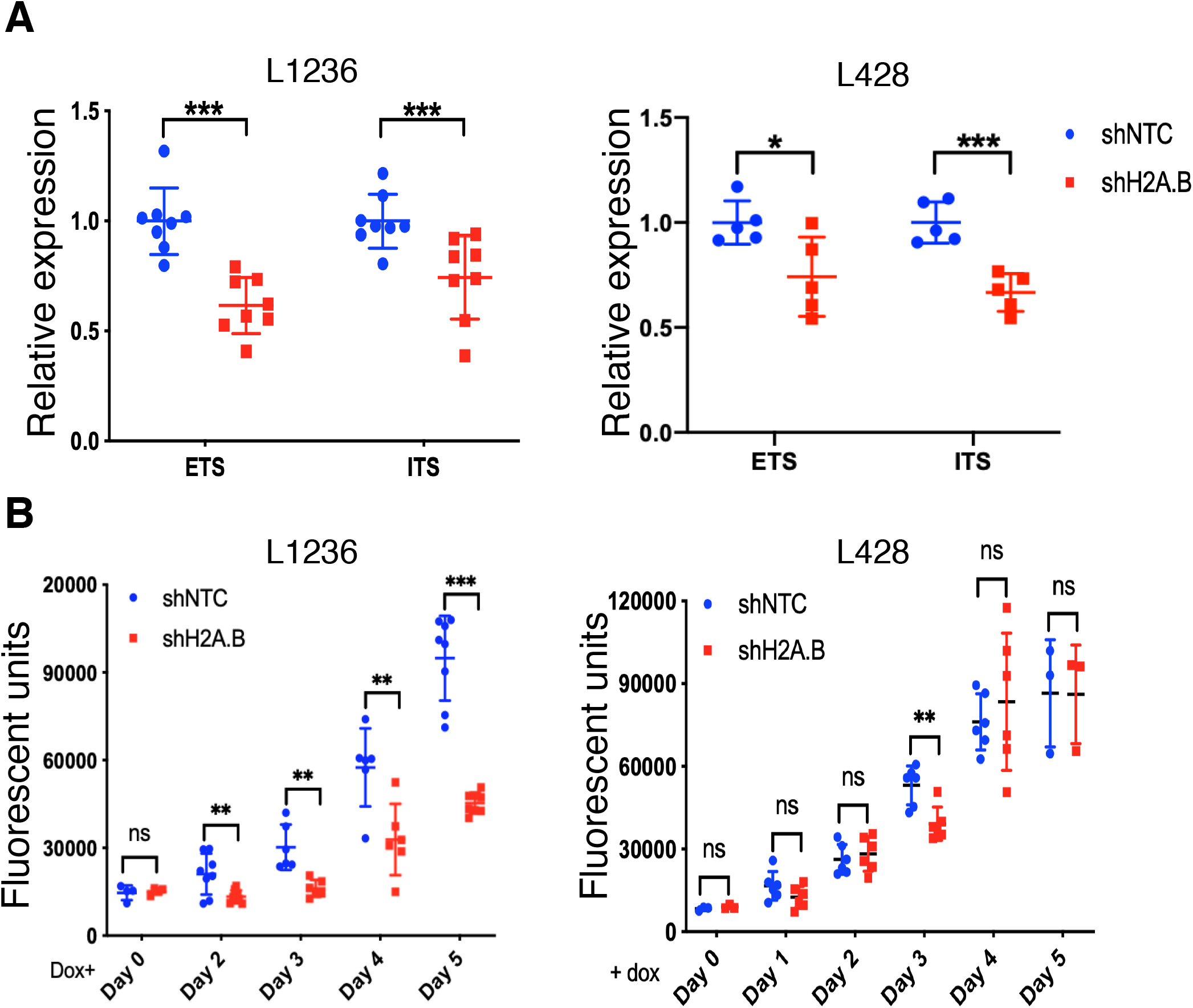
H2A.B is required for high levels of pre-rRNA transcription. (**A**) qPCR of cDNA from L1236 and L428 HL cell RNA obtained 5 days post H2A.B knockdown. Amplification was performed using primers that anneal to the ETS and ITS spliced-out regions to detect nascent rRNA. ETS and ITS, external and internal transcribed regions, respectively. (**B**) Cell proliferation assay in the presence (shNTC) or during depletion of H2A.B (shH2A.B). The results are presented as mean ± SD; Student’s t test, ns, p > 0.05; *, p<0.05; **, p < 0.01; ***, p<0.001 (n=5).

High levels of rRNA transcription have been shown to have a direct link to high rates of cell proliferation in cancer (Ferreira et al., 2020). Given that H2A.B enhances rDNA transcription, we next assessed whether the loss of H2A.B can affect the rate of HL cell line proliferation. Cell proliferation assays indeed showed that H2A.B knockdown caused a marked reduction in the proliferation rate in L1236 HL cells (Figure 4B). Although not as significant, a similar trend was observed in L428 HL cells. This and other differences observed between L1236 and L428 HL cells can be attributed to the less efficient knock down of H2A.B expression in L428 compared to L1236 cells (in L428 HL cells, ~25% of H2A.B protein still remains).

### Depletion of H2A.B downregulates the expression of ribosomal protein genes

Given that H2A.B facilitates RNA Pol II transcription in the mouse testis (Anuar et al., 2019; Soboleva et al., 2012; Soboleva et al., 2017), we next sought to investigate the impact of H2A.B knockdown on RNA Pol II-driven gene expression in the L1236 and L428 HL cell lines (three biological replicates on day 5 post-induction of shH2A.B versus shNTC RNA expression). We found that 4,775 genes (36.5%) had a significant change in expression (FDR≤ 5%) in L1236 HL cells: 48.5% were downregulated and 51.5% were upregulated (Figure 5A). Similarly, for L428 HL cells, 3,937 genes (29.4%) had a significant change in expression pattern: 2,108 (53.5%) genes were downregulated and 1,829 (46.5%) were upregulated (Figure 5B).

**Figure 5.**
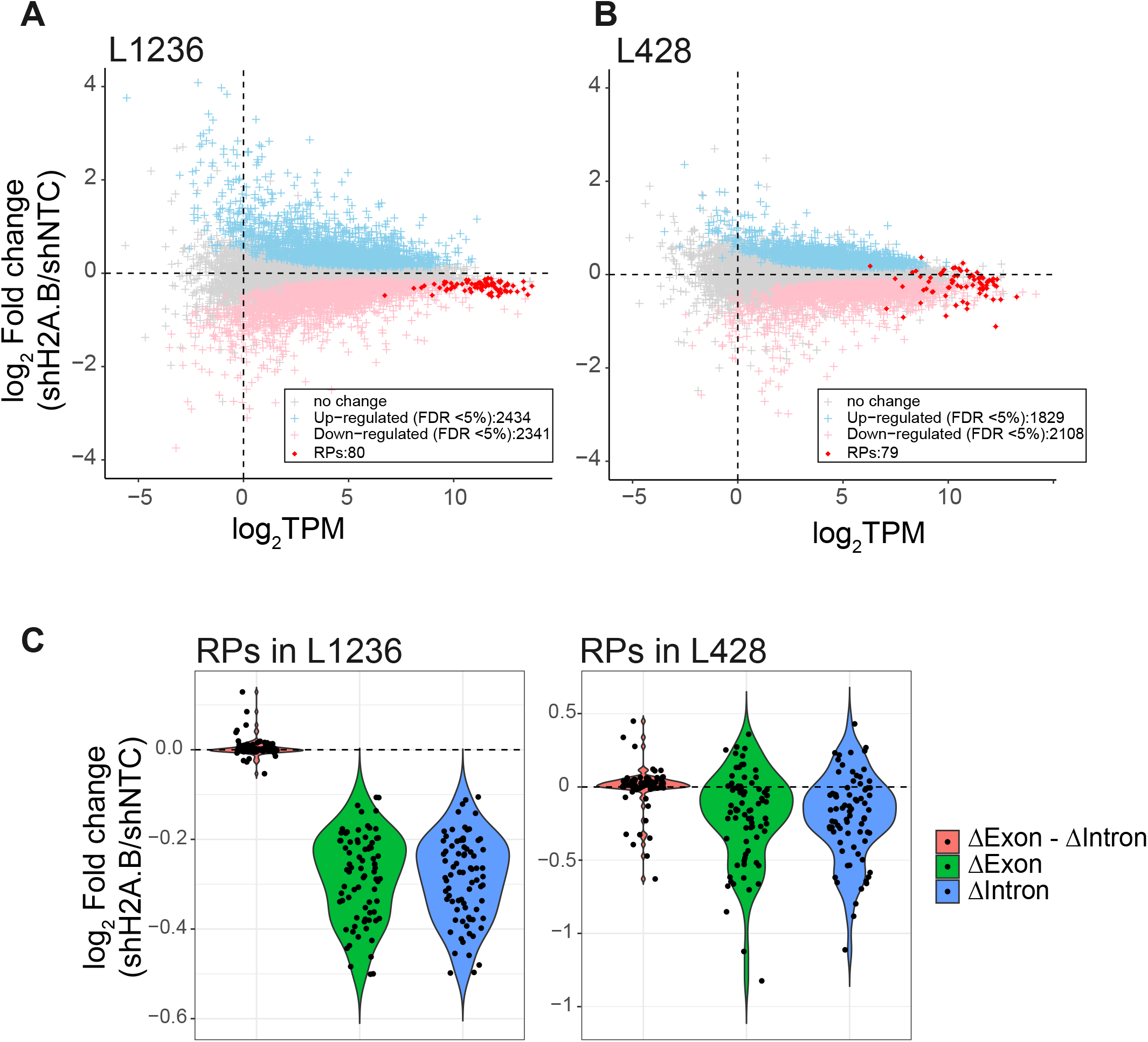
H2A.B is required for high levels of ribosomal protein transcription. Differentially expressed genes between shNTC and shH2A.B in **(A**) L1236 HL cells and **(B**) L428 HL cells. Gene expression fold changes (Log2) are plotted against the gene expression level (Log2 TPM). For both cell lines, RP genes are highlighted in red. (**C**) The RNA-seq data were analysed by Exon-Intron Split Analysis. RNA-seq reads were separated into exonic (Δexon) and intronic (Δintron) changes. Exonic changes (Δexon) reflect the combination of changes in transcriptional and post-transcriptional activity changes. Intronic changes (Δintron) reflect changes in transcriptional activity. Δexon – Δintron reflects post-transcriptional effects. RP genes in L1236 and L428 HL cells are shown.

Most interestingly, the differentially expressed genes were almost equally spread between upregulated and downregulated (Figure 5A and B), but this trend changed for the most highly expressed genes. For genes whose log_2_TPM was ≥ 10, expression was almost exclusively downregulated following H2A.B knockdown in both cell lines. This finding implicates a role of H2A.B in sustaining high levels of transcription for the most highly expressed genes. Notably, ribosomal protein (RP) genes were among the most highly expressed genes. We found that 56/80 of RP genes were significantly downregulated in L1236 cells and that 28/79 of RP genes were significantly downregulated in L428 cells (FDR≤ 5%; Figure 5A and B; RP genes highlighted as red dots).

This decrease in RP gene expression was validated in L1236 cells by qPCR analysis of 26 RP genes, which included 14 large ribosomal subunit (RPL) and 12 small ribosomal subunit (RPS) genes (Supplementary Figure 6). Consistent with the RNA-seq data, 24 of these 26 genes showed a significant decrease in gene expression following H2A.B knockdown. In combination, it can be concluded that H2A.B has a major role in ensuring high levels of RP gene expression.

RP gene expression is known to be subject to post-transcriptional regulation, including alterations to their mRNA stability (Gentilella et al., 2017). Therefore, the RNA-seq data were analysed further by Exon-Intron Split Analysis (EISA) (Gaidatzis et al., 2015). This analysis differentiates between transcriptional and post-transcriptional changes by comparing the changes in exonic (mature mRNA) and intronic (pre-mRNA) reads for the same gene. With an FDR < 5%, 74/77 RPs show significant exonic (Dexon) and intronic (Dintron) changes. In contrast, no significant post-transcriptional change (Δexon – Δintron) was observed for any RP gene (as shown by a Log2shH2A.B/shNTC of around zero; Figure 5C). This observation indicates that the decrease in RP gene expression can be attributed to a reduction in the level of transcription. The same trend was observed for L428 HL cells (Figure 5C).

### H2A.B-containing nucleosomes are enriched at the TSS and gene body of RP genes

Previously, we have shown that H2A.B is targeted to the TSS and gene body of Pol II genes, including intron–exon boundaries, concurrent with their activation in the mouse testis (Anuar et al., 2019; Soboleva et al., 2012; Soboleva et al., 2017). To investigate whether H2A.B occupies the same genic locations in the HL cancer setting, we performed Cleavage Under Targets and Release Using Nuclease (CUT&RUN) experiments (Skene et al., 2018). This method is an alternative to ChIP-seq. It relies on antibody–MNase targeted cleavage of native chromatin within a permeabilized cell nucleus followed by paired-end sequencing of the released DNA fragments. This method has the significant advantage of reducing background reads. A metagene H2A.B CUT&RUN analysis for all RP genes, from 1 kb upstream of the transcriptional start site to 1kb downstream of the transcription termination site, showed high occupancy of a H2A.B-containing nucleosome positioned at the TSS and that these variant nucleosomes extended into the gene body in L1236 and L428 HL cells (Figure 6A and B, respectively). Combined with the inhibition of RP gene expression following H2A.B knockdown, this shows that H2A.B is directly responsible for the high level of RP gene transcription.

**Figure 6.**
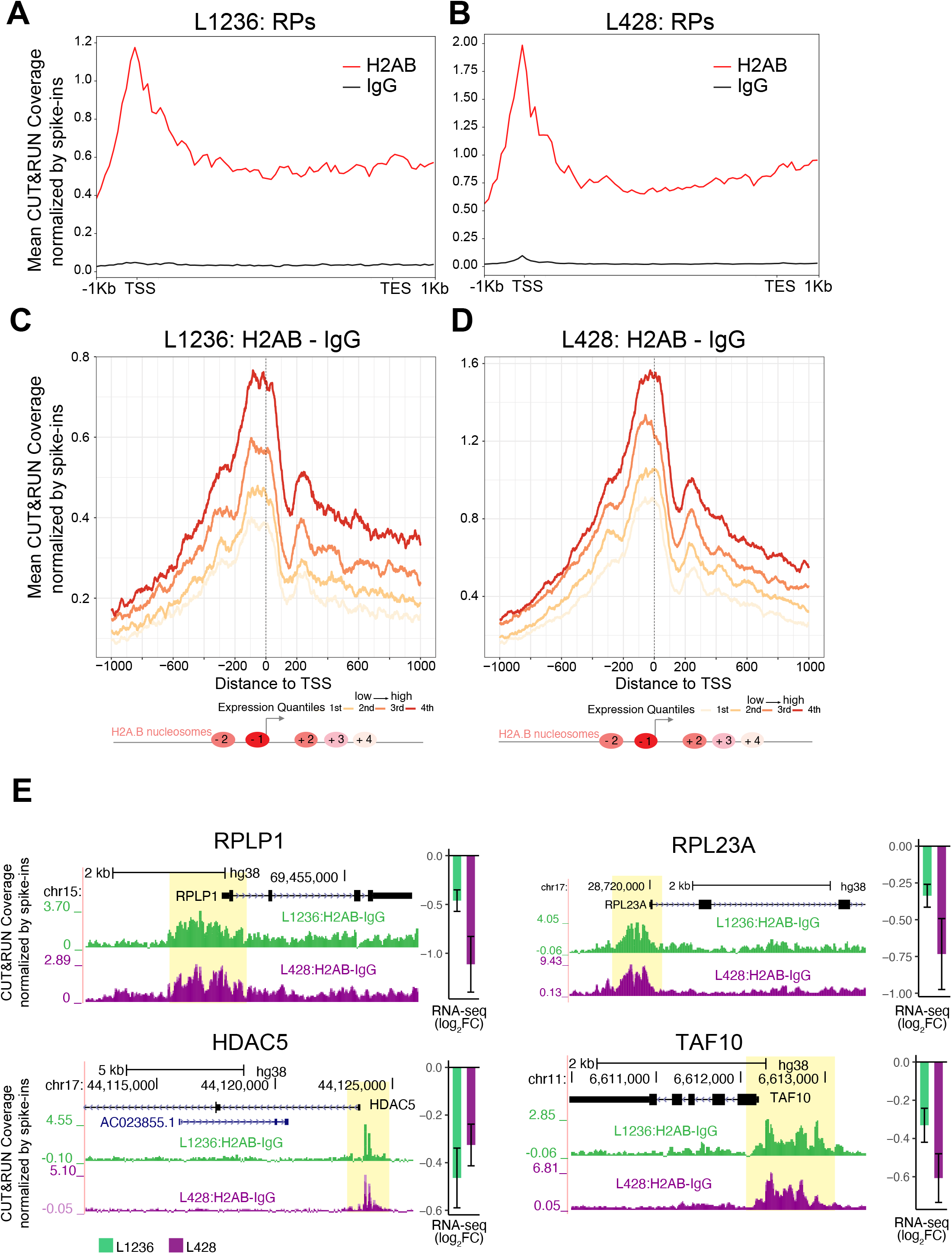
H2A.B is located at the TSS of active genes. Mean H2A.B and IgG CUT&RUN coverage normalised by spike-ins from 1kb upstream to 1kb downstream for all ribosomal protein genes in **(A**) L1236 HL cells and **(B**) L428 HL cells. Mean difference between H2A.B and IgG CUT&RUN coverage normalized by spike-ins aligned between −1 and +1 kb from the TSS ranked according to their level of expression in **(C**) L1236 HL cells and **(D**) L428 HL cells. Below is the position of H2A.B nucleosomes relative to the TSS; the colour reflects nucleosome abundance (red reflects the highest abundant H2A.B nucleosome located at the TSS). (**E**) Genome browser screen shots of gene examples (RPs RPLP1 and RPL23A, and HDAC5 and TAF10) that contain H2A.B at their promoters identified by the CUT&RUN peak calling and whose expression decreased significantly following H2A.B knockdown in the RNA-seq analysis. Green and purple indicates L1236 and L428cells, respectively.

To confirm that all active Pol II genes display this H2A.B genic localization pattern, genes were separated into quartiles according to their expression level and classified as repressed, low, medium or high. For each of these groups of genes, a single line represents H2A.B reads at each base pair aligned with the TSS (±1 kb). As observed in the mouse testis (Soboleva et al., 2012; Soboleva et al., 2017), a strongly positioned H2A.B-containing nucleosome was centred at ~ 50 bases upstream of the TSS, and its appearance correlated positively with gene expression in both HL cell lines (Figure 6C and D). It is also found downstream of the TSS but is excluded from the important +1 nucleosome. The RPs RPL1 and RPL23A, the histone deacetylase HDAC5, which has been shown to have an important role in cancer (Cao et al., 2017), and TAF10, the component of the transcription initiation factor TFIID, are examples of active genes that contain H2A.B at their TSSs. Also as seen in the mouse testis (Soboleva et al., 2017), H2A.B was targeted to exon–intron (and intron–exon) boundaries, and its abundance at these sites of splicing is also correlated positively with transcription (Supplementary Figure 7).

Next, we investigated the overlap of H2A.B genomic locations between the two different cell lines. Peak calling of H2A.B.CUT&RUN data revealed that, for both cell lines, the majority of H2A.B peaks were localized within intronic regions (~60%), followed by promoter-TSS regions (~10%) and intergenic regions (~10%) (Supplementary Figure 8A). Analysis of the peak overlap between L1236 and L428 HL cell lines revealed that the majority of peaks overlap in the promoter–TSS region compared with other genomic locations. About 50% and ~60 % of L1236 and L428 peaks overlapped in the promoter-TSS region, respectively compared with ~17% and ~29% for L1236 and L428 peaks for all other genomic locations, respectively (Supplementary Figure 8B). Given this strong overlap of H2A.B at promoters between the two HL cell lines, this finding implies that H2A.B regulates the transcription of common genes and gene expression pathways. Therefore, we next investigated the gene expression pathways and networks that are commonly regulated by H2A.B in both cell lines.

### H2A.B regulates pathways involved in post-translational modification and HIF-1

We found that 812 genes were commonly downregulated in response to H2A.B knockdown in L1236 and L428 HL cell lines and that 404 of these genes contained H2A.B peaks within the promoter-TSS region (± 1Kb from the TSS; Supplementary Figure 8C and D, Supplementary Table 2). A Uniprot keyword enrichment analysis revealed that these genes are highly associated with terms that represent post-translational modification pathways that include acetylation, phosphorylation and ubiquitylation, as well as the expected ribosomal and ribonucleoprotein pathways (Supplementary Figure 8E, Supplementary Table 3). A KEGG pathway analysis also revealed the ribosome as the major pathway regulated by H2A.B in both cell lines (Supplementary Figure 8E, Supplementary Table 3; see this table also for Reactome and GO analyses).

Interestingly, the 161 proteins in the acetylation network (Supplementary Table 4) not only includes enzymes that synthesize acetyl-CoA (ATP-citrate synthase) and deacetylate histones (notably HDAC5, as noted above), but also contain other histone modifying enzymes (methyltransferases and histone demethylases). TAF10 is also part of this acetylation network as are RPs, an observation that further strengthens the link between H2A.B and ribosome biogenesis. Taken together, these findings suggest that the impact of H2A.B on chromatin function may go beyond the direct regulation of gene expression but may also affect chromatin function and transcription indirectly by influencing post-translational modifications.

The KEGG pathway analysis also revealed that the transcription factor hypoxia-inducible factor-1 (HIF-1) pathway is downregulated in both cell lines in response to H2A.B KD (Supplementary Figure 8E). HIF is a major regulator of hypoxic responces and consequently activates ID2, NOTCH1, AP-1, NFκB and JAK/STAT signalling pathways, which are all hallmarks of HRS cells (Kuppers et al., 2012). There is evidence that the activation of HIF-1 signaling during the early stages of HL development may lead to de-differentiation of the B-cell phenotype and its reprogramming into HRS cells (Wein et al., 2015). This data raises the intriguing possibility that H2A.B may contribute to the establishment the HRS phenotype by regulating the HIF-1 pathway.

### H2A.B regulates pre-mRNA splicing

To address whether H2A.B has a role in regulating pre-mRNA splicing in HL cells, we examined the exon usage in H2A.B knockdown versus control L1236 cells at day 5 post-induction. Significantly, the loss of H2A.B impaired pre-mRNA splicing, which caused an overall increase in the inclusion of alternatively spliced cassette exons in L1236 HL cells (Figure 7A). We observed 2,374 genes to have significantly (FDR < 10%) differential exon usage after H2A.B knockdown, and this included a total of 4,677 differentially spliced exons. *CD44* is an example of a gene that displayed such altered splicing events (Figure 7B). *CD44* is linked to metastasis and distinct spliced isoforms are common in cancer (Sveen et al., 2016). Strikingly, splicing of 68% (54/80) of RP transcripts was also altered as illustrated for RPL17 (Figure 7C). This indicates that the loss of H2A.B not only affects the level of RP transcription but may also regulate the synthesis of different RP transcript isoforms. Similar changes in splicing were also observed in L428 HL cells, as shown for TIMM44 (mitochondrial inner membrane translocase), KNOP1 (lysine-rich nucleolar protein 1), PI4K2B (phosphatidylinositol 4-kinase type II beta), and the RP gene RPS16 (Supplementary Figure S9). Therefore, the role of H2A.B in HL is multi-layered affecting both transcription and splicing.

**Figure 7.**
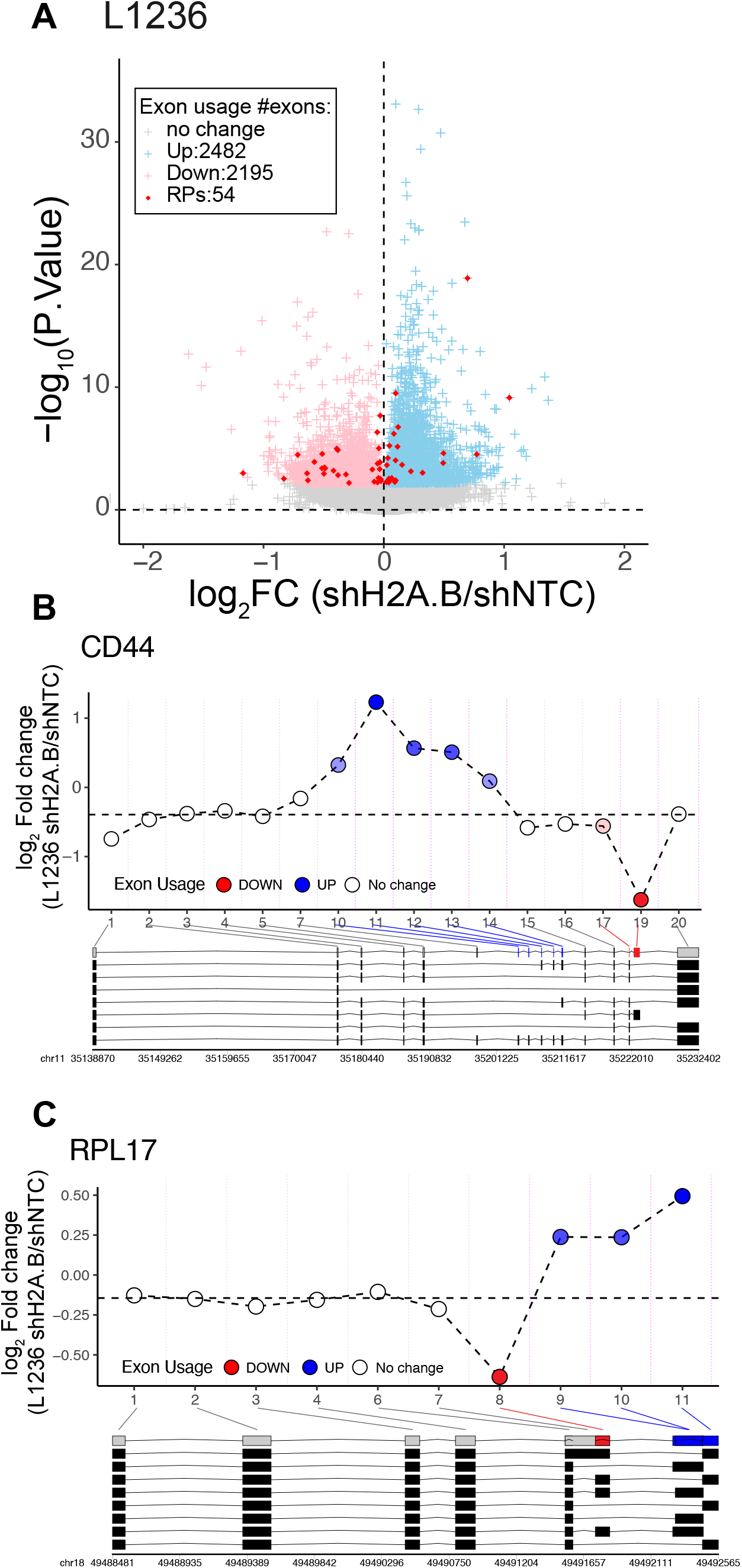
Pre-mRNA splicing is altered following the depletion of H2A.B. (**A**) Changes in exon usage between shNTC and shH2A.B L1236 HL cells. Highlighted in red are RP genes. The significance (-log_10_ (P value)) is plotted against the fold change of exon usage (log_2_FC). (**B**) Differential exon usage of the CD44 gene. (**C**) Differential exon usage of the RPL17 gene.

### H2A.B modulates chromatin accessibility

*In vitro* chromatin reconstitution and transcription assays have demonstrated that the incorporation of H2A.B decompacts chromatin thereby enabling DNA access to the transcription machinery (Angelov et al., 2004; Soboleva et al., 2012; Zhou et al., 2007). We therefore sought to determine whether the loss of H2A.B would alter chromatin accessibility in cells. To investigate this, we used the Transposase-Accessible Chromatin followed by sequencing (ATAC-seq) assay on L1236 cells 5 days post-H2A.B knockdown and compared accessibility with that of the non-targeting shRNA control. A meta-gene plot of accessibility across the intron-exon boundary for the top quartile of expressed genes (Supplementary Figure 10A) and the highly transcribed RP genes (Supplementary Figure S10B) revealed a decrease in accessibility following the knockdown of H2A.B expression.

To further explore the relationship between chromatin accessibility and H2A.B more rigorously, the change in accessibility of all H2A.B CUT&RUN peaks was examined in H2A.B knockdown L1236 HL cells. H2A.B knockdown produced 5,552 peaks with significantly increased accessibility and 3,310 peaks with significantly decreased accessibility (FDR≤10%) (Figure 8A). However, the extent of decreased accessibility was more pronounced than that of increased accessibility (Figure 8B). Investigation of the genomic location where these changes in accessibility occurred produced striking results. The increase in accessibility occurred mainly at promoters (40% of H2A.B peaks such as the NXF1 promoter) (Figure 8C and D), whereas the decrease in accessibility following the depletion of H2A.B occurred mainly in introns (70% of H2A.B peaks such as a KDM4A intron) (Figure 8C and E).

**Figure 8.**
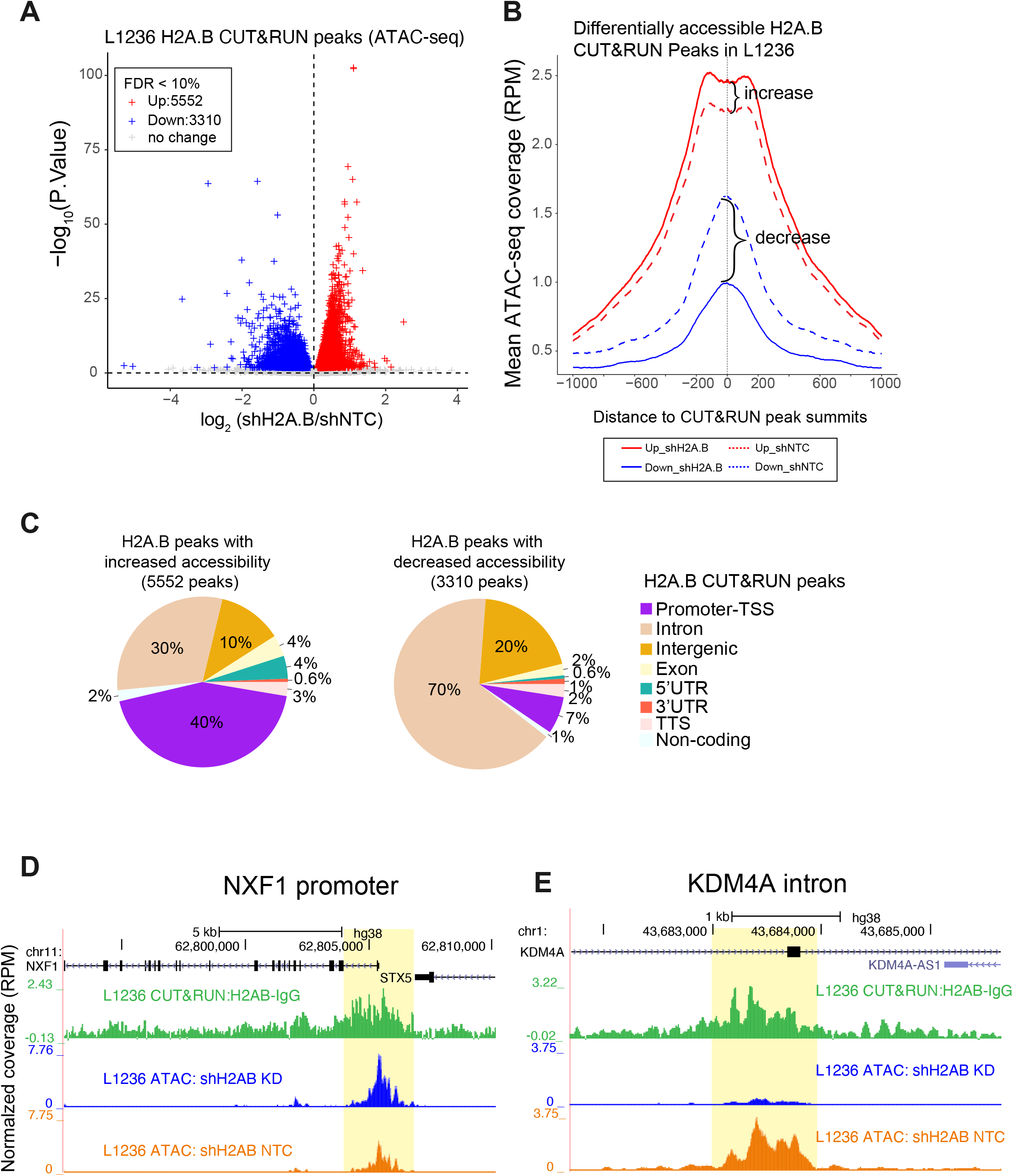
Loss of H2A.B decreases chromatin accessibility in introns. (**A**) Changes in ATAC-seq accessibility at H2A.B CUT&RUN peaks following H2A.B knockdown. The significance (-log_10_ (P value)) is plotted against the fold change in accessibility (log_2_ (shH2A.B/shNTC)). Red indicates H2A.B peaks that increased in accessibility (5,552). Blue indicates H2A.B peaks that decreased in accessibility (3,310). (**B**) Differential accessibility changes of H2A.B CUT&RUN peaks following H2A.B knockdown for the peaks that increased in accessibility (red) and peaks that decreased in accessibility (blue) in Panel A. The change in mean ATAC-seq coverage (RPM) is plotted against the centre of the H2A.B peaks. (**C**) Genomic distribution of H2A.B CUT&RUN peaks, shown as percentages, with increased (5,552 in total) or decreased (3,310 in total) accessibility following H2A.B knockdown. Two examples show (**D**) increased accessibility at the promoter peak (NXF1) and (**E**) decreased accessibility at the peak across the intron–exon boundary (KDM4A), respectively.

These results show that there is a major decrease in chromatin accessibility within the gene body (in agreement with the data presented in Supplementary Figure 10). These findings are consistent with the notion that chromatin becomes more compact in the absence of H2A.B. By contrast, the promoter–TSS becomes slightly more accessibility perhaps because the TSS now becomes nucleosome free.

## DISCUSSION

Although the first C/T gene, MAGE-1, was discovered almost 30 years ago (van der Bruggen et al., 1991), and to date more than a thousand putative C/T genes have been identified (da Silva et al., 2017; Wang et al., 2016), surprisingly little is known about the molecular function of C/T genes in cancer. Further, a feature of virtually all types of cancer is the major alterations to the epigenome, which contributes to abnormal patterns of transcription and splicing (Baylin and Jones, 2011; Rousseaux and Khochbin, 2009; Wang et al., 2011), and a subset of C/T genes expressed in cancer encode proteins that are modifiers of the epigenome (Debruyne et al., 2019; Simpson et al., 2005). However, the role of such testis epigenome regulators in establishing the cancer phenotype is poorly understood because only a small number of such C/T genes have been studied in detail. Elucidating their mechanism of action may provide new avenues for the treatment of cancers.

For example, the C/T gene BORIS, a germ-cell-specific paralogue of the chromatin architectural protein CTCF, is upregulated in neuroblastoma cells (Debruyne et al., 2019). BORIS promotes new chromatin interactions leading to the formation of super-enhancers, which drives the expression of a group of transcription factors rendering these cancer cells resistant to treatment. BRDT is another CT gene that regulates the epigenome. BRDT is a member of the double bromodomain BET family and is aberrantly activated in lung and breast cancers (Bourova-Flin et al., 2017). BRDT binds to hyperacetylated histone H4, compacts chromatin, and is required for the exchange of histones with protamines during spermatogenesis. It is also required for the establishment of meiotic and post-meiotic gene expression programs by recruiting Positive Transcription Elongation Factor (P-TEFb) (Gaucher et al., 2012). It is proposed that these functions of BRDT contribute to malignant transformation when it is aberrantly expressed (Bourova-Flin et al., 2017).

The data presented here show that the ectopic expression of H2A.B is a common feature of primary HL because it was detected in every tumour analysed independent of the HL subtype. This provided us with the basis to test the hypothesis that the expression of testis regulators of chromatin function can play an important role in promoting oncogenesis by studying the role of H2A.B in HL derived cell lines. Further, to test this hypothesis rigorously and to obtain insights into the HL gene expression pathways regulated by H2A.B, we analysed and compared two different HL cell lines that were originally derived from different HL subtypes.

One characteristic feature of HRS is their large and highly polymorphic nucleoli, which are reflective of high levels of rDNA transcription (Mamaev et al., 1997). A major finding of this study is that H2A.B enhances ribosome biogenesis by two different mechanisms. First, it elevates the level of Pol I transcription. This is a cancer acquired function because such a role is not seen in the mouse testis. Therefore, histone variants can assume new functions when expressed in a non-physiological context.

Second, H2A.B increases the transcription of RP genes. An increase in ribosome biogenesis is a common feature of cancer cells as it is required to sustain high growth rates (Drygin et al., 2011; Ferreira et al., 2020). Moreover, blocking the production of new ribosomes inhibits cell proliferation (Pestov et al., 2001; Volarevic et al., 2000). Consistent with a role of H2A.B in ribosome biogenesis, knocking down its expression caused a significant reduction in the rate of HL cell proliferation.

Several studies have shown that when H2A.B is ectopically expressed; it accumulates in the nucleolus (Ioudinkova et al., 2012; Shaw et al., 2009). However, it was unclear whether H2A.B had a role in nucleolar function or if over-expressed H2A.B simply accumulates in the nucleolus to be ultimately degraded. Here, we have demonstrated for the first time a role for H2A.B in Pol I transcription by showing that: 1) H2A.B is enriched in the nucleolus of HRS cells; 2) it is located at the rDNA promoter and transcribed regions; 3) H2A.B directly interacts with RNA Pol I; and 4) the knockdown of H2A.B expression reduces the level of Pol I at rDNA chromatin with a corresponding decrease in the level of rDNA transcription. In establishing a role for H2A.B in RP Pol II transcription, we demonstrate that: 1) H2A.B-containing nucleosomes are found at the TSS and in the gene body, including intron–exon boundaries of highly expressed RP genes and 2) the knockdown of H2A.B expression reduces the level of transcription of a large number of RP genes. Further, in mouse spermatids, H2A.B interacts with RNA Pol II (Soboleva et al., 2017). Taken together, these findings indicate that H2A.B can increase the production of mature ribosomal subunits by enhancing both rRNA and ribosomal protein synthesis.

The expression of RPs can be controlled at the level of splicing to produce different protein isoforms (Ivanov et al., 2006). Significantly, we also revealed that H2A.B can regulate the pre-mRNA splicing of RPs suggesting that H2A.B can regulate ribosome biogenesis both quantitatively and qualitatively.

Another major finding of this study is that the contribution of H2A.B to the HL phenotype appears to be multifaceted by extending beyond its regulation of ribosome biogenesis. H2A.B regulates the expression and splicing of many other Pol II transcribed genes and some of these are implicated in cancer progression, such as CD44 (Sveen et al., 2016), HDAC5 (Cao et al., 2017) and HIF1 (Wein et al., 2015). Further, we identified the common genes between L1236 and L428 HL cell lines whose expression is stimulated by H2A.B. Unexpectedly, many of these genes are in pathways that involve post-translational modifications, which suggests that H2A.B may regulate the function of many other proteins indirectly. For example, in the acetylation network, the expression of HDAC5 in HL cells is controlled by H2A.B. Other genes in this network include components of the basal transcription factor TFIID (TAF10), histone methyl transferases and demethylases (CARM1 and KDM4A) and transcription factors (e.g. MAX and ARID3B). Therefore, the impact of this single histone variant on HL nuclear function appears to be substantial.

From a structural perspective, H2A.B (and its family member H2A.L.2; (Barral et al., 2017)) is a unique histone variant because of its ability to inhibit intra-nucleosome-nucleosome interactions and chromatin compaction to overcome chromatin-mediated repression of transcription *in vitro* (Angelov et al., 2004; Soboleva et al., 2012; Zhou et al., 2007). However, to our knowledge, no study has investigated whether H2A.B can regulate chromatin compaction and accessibility in cells. Upon the knockdown of H2A.B, chromatin accessibility within the body of genes (at introns) decreases significantly which is consistent with chromatin becoming more compact. At a more local level, promoter–TSS regions became slightly more accessible perhaps because the TSS now becomes nucleosome free. We propose that the ability of H2A.B to decompact chromatin plays an important role in increasing transcription in HL cells.

Analogous to H2A.B, the testis specific histone variant dimer pair TH2A and TH2B are also able to decompact chromatin and increase chromatin accessibility (Hoghoughi et al., 2018). Most interestingly, it was reported that expression of TH2A/TH2B together with the Yamanaka transcription factors helped to reprogram somatic cells into iPS cells more efficiently than Yamanaka factors alone (Shinagawa et al., 2014). Future work will determine whether the ectopic expression of H2A.B also makes the somatic cell genome more responsive to reprogramming.

In conclusion, this study provides strong support for the hypothesis that the unique functions and properties of H2A.B have been hijacked by HL to transform the nucleolus to a more active state, and to reprograme the epigenome to alter patterns of transcription and splicing. This demonstration that H2AFB1 is a cancer/testis gene may provide alternative avenues for the diagnosis and/or treatment of HL.

## MATERIALS AND METHODS

### Cell lines

HL cell lines L1236 and L428 were cultured in RPMI-1640 growth medium supplemented with 10% heat-inactivated foetal bovine serum (HI-FBS), 1% L-glutamine (200 mM stock), and 1% Penicillin–Streptomycin (10,000 units penicillin and 10 mg streptomycin/ml stock). Human embryonic kidney HEK293T cells were cultured in Dulbecco’s modified Eagle’s medium (DMEM) supplemented with 10% HI-FBS, 1% L-glutamine, and 1% Penicillin-Streptomycin. Cells were cultured in a humidified cell culture incubator at 37°C and 5% CO2.

### Antibodies

H2A.B (rabbit polyclonal) WB, 1: 1000; IHC 1:100; CUT&RUN 1: 50; IP 1: 250; β-actin (8457S, Cell Signalling) WB, 1:10,000; RPA194(sc-48385, Santa Cruz) WB, 1: 1000; ChIP, 10 μl/40 μg DNA; H3K3me36 (used as a positive control for CUT&RUN, ab9050, Abcam) CUT&RUN 1:200; CD15 (GA062, Dako Omnis) IHC 1:100; Fibrillarin (Ab4566, Abcam) IHC/IF 1:100; H3 histone (Ab1791, Abcam) WB 1:250,000; V5-FITC (Ab1274, Abcam) IF 1:200; anti-rabbit IgG-HRP (111-035-144, Jackson Immuno), WB1:50,000; Anti-rabbit-IgG-Cy3 (713-165-152, Jackson Immuno) IF/IHC 1:200; antimouse IgG-FITC (715-095-150, Jackson Immuno) IF/IHC 1:100.

### Nucleic acid extraction, RT-PCR, and qPCR

Total RNA was extracted from 1×10^8^ cells using TRIzol (Thermo Fisher) reagent. The RNA was treated with DNaseTURBO (Thermo Fisher) before qPCR or RNA-seq library preparation. cDNA was prepared from 1 μg of total pre-treated RNA with SuperScript™ III (Thermo Fisher) and an equimolar ratio of Oligo-dT and random hexamer primers. For qPCR, 0.5–2 μl of DNA template was used in a 10 μl reaction with gene-specific primers. *B2M* and *HPRT1* served as endogenous reference controls for measuring RNA expression. The results were calculated using the 2^-ΔΔCT^ method (Rao et al., 2013). For ChIP–qPCR, the enrichment level in the target locus was normalized to the input, and the results are presented as fold changes relative to the enrichment level at the enhancer repeat element (ENHCR) site. For genomic DNA extraction, ans Isolate II Genomic DNA kit (Bioline) was used to extract genomic DNA from 1×10^7^ cells. For Sanger sequencing, 10 ng of DNA was used with primers for H2AFB1-3 in PCR reaction. All primer sequences are listed in Supplementary Table 1.

### Expression and purification of recombinant H2A.B

Codon-optimised H2AFB3 gene DNA sequences flanked with *Bam*HI and *NdeI* restriction sites, was synthesized by Life Technologies™ and cloned into pET-3a expression vectors (Novagen). Rosetta2 (DE3) pLysS *E. coli* cells (Novagen) were used for protein expression using 1 L of auto-induction media (1 % tryptone, 0.5 % yeast extract, 0.5 % glycerol, 0.05 % glucose, 0.2 % α-lactose, 1 mM MgSO4, 100mM (NH_4_)_2_SO_4_, 50 mM KH_2_PO_4_, 50 mM Na_2_HPO_4_) overnight at 37°C. Following autoinduction, inclusion bodies were purified following a standard protocol. Next, inclusion bodies were dissolved in unfolding buffer (6 M Guanidine HCl, 20 mM Na Acetate, 1 mM DTT, pH 5.2) and dialysed overnight against 1L SAUDE 600 buffer (7 M Urea, 20 mM Na Acetate, 600 mM NaCl, 5 mM b-mercaptoethanol, 1 mM EDTA, pH 5.2). The dialysate was separated using Superdex 200 gel filtration column. The H2A.B protein was eluted and fractions containing pure H2A.B were pooled together and dialysed overnight against two changes of 2.5 % acetic acid and 5 mM b-mercaptoethanol. After dialysis, the protein was aliquoted and lyophilized using Dura-Dry™ microprocessor control corrosion resistant freeze-dryer (Kinetics).

### Anti-H2A.B antibody production and purification

Antigen injections were performed by SAHMRI Preclinical, Imaging and Research Laboratories (PIRL) in Adelaide, Australia. Two rabbits were used for the antibody generation. Each rabbit was immunized by four injections of the 0.25 μg of purified recombinant human H2A.B at three-week intervals. For affinity purification of the H2A.B-specific IgGs from crude serum samples, the recombinant H2A.B protein immobilized on a PVDF membrane was used as the antigen.

### Primary Hodgkin Lymphoma tumour sample selection

Formalin-fixed paraffin-embedded postsurgical lymph node specimens were retrieved from the archival material in ACT Pathology, the Canberra Hospital, following ethics approval by the ACT Health Human Research Ethics Committee (ETHLR 14.260). This included blocks from 81 randomly selected deidentified patients with Hodgkin’s lymphoma diagnosed between 1997 and 2014. Their initial diagnosis was confirmed on review of the light microscopy and immunohistochemistry. The age and gender of the patients was provided but no prognostic data was available. Out of 81 cases, 51 patients were diagnosed with nodular sclerosing HL, 14 mixed cellularity HL, 4 cases with lymphocyte rich HL, 6 cases of nodular lymphocyte predominant HL and 6 not otherwise classified. There were no cases of lymphocyte depleted HL in this cohort.

### Immunohistochemistry of primary HL samples

Tissue microarrays (TMAs) were constructed with four cases per slide. A positive control (adult testis) and a negative control (tonsil tissue) were included on each slide. The immunohistochemistry (IHC) staining was firstly optimized by using round spermatids in human testis and lymphocytes in tonsil tissue. Immunohistochemistry was performed on the Ventana automated system (BenchMark ULTRA), following a standard protocol. Briefly, tissue sections were cut (3μm thick) and dried in an oven at 60°C for 20 min. Heat retrieval was performed for 8 min followed by blocking for 24 min. The sections were stained with 1:1000 dilution of H2A.B rabbit polyclonal antibody incubated for 32 min. UltraView DAB IHC Detection Kit (Ventana) was used to amplify and visualize the signals. HRS cells were visualised by co-staining with CD30 antibodies. Nucleolar localisation of H2A.B was determined by co-staining with Fibrillarin. Nuclei were visualized by counterstaining with Haematoxylin. H2A.B signal was scored for each sample, for HRS cells and infiltrating lymphocytes separately, at least 4 fields of approximately 100 cells of each field were analysed.

### Establishment of inducible H2A.B knockdown HL cell lines by lentiviral transduction

HEK293T at 40% confluency were transfected with 20 μg of SMARTvector (containing TurboRFP and Puro^r^ genes for selection; Tet-ON-3G for doxycycline (dox) induction, Dharmacon) coding shH2A.B (ATTGAGTACCTGACGGCCA) or shH2A.B-2 (CCAGGTGGAGCGCAGTCTA) or shNTC (proprietary sequence of Dharmacon), respectively; 15 μg psPAX2 packaging plasmid; 6 μg pMD2.G envelope expressing plasmid and 0.4M CaCl_2_ in a HEBS buffer (25mM HEPES -NaOH, 140mM NaCl, 5 mM KCl, 0.75mM Na2HPO4, and 6mM glucose, pH 7.05). 16-24 h post-transfection, the cell medium was replaced, incubated for further 6-9 h, replaced again and incubated for 16-24 h followed by collection of virus-containing media. For transduction of L1236 and L428 cells, viral media was supplemented by 10 μM

HEPES-NaOH pH 7.4 and 4 μg/ml polybrene. The 2-4 x10^6^ of target cells were transduced by spinoculation at 2,000 RPM for 1 h at room temperature. The post-transduced cells were selected with 0.4 μg/ml puromycin or 0.5 mg/ml G-418 (for L1236) or with 4 μg/ml puromycin or 1 mg/ml G-4186 days (for L428) for 6 d. The L428 transduced cells were further selected by FACS based on the level of TurboRFP expression, since a large portion of the L428 cells produce relatively weak TurboRFP signal after antibiotic selection. The cells were induced by 1 μg/ml doxycycline for 48 h before sorting. Finally, shRNA expression was induced by addition of dox at 0.1 μg/ml for L1236, and 1 μg/ml dox in L428 cells for up to 5 days to achieve levels of down-regulation of H2A.B greater than 70-80%.

### Establishment of exogenous H2A.B-expressing cell lines

pEF1 *a-V5-H2A.B-IRES-P* construct was transiently transfected in 70% confluent HEK293T cells using 1 μg of plasmid DNA and 2 μl of Lipofectamine™2000 per well in the 24-well plates. The immunofluorescent staining was performed 24 h posttransfection using V5-FITC (Abcam) and purified anti-H2A.B antibody.

For proximity biotin labelling stable L428 and L1236 cell lines were created overexpressing two constructs: either BirA* (R118G) at the N-terminus (V5-BirA*-H2A.B) or the C-terminus (H2A.B-BirA*-HA).The H2A.B-BirA*-HA was created by cloning H2A.B into pLVX-IRES-ZsGreen1 vector using EcoRI and XbaI restriction sites, then adding BirA*-HA from pcDNA3.1 MCS-BirA(R118G)-HA vector. The H2A.B-BirA*-HA construct was then cloned into pLVX-EF1a-IRES-Neo vector using the EcoRI and BamH1 restriction sites, giving rise to the pLVX-EF1a-H2A.B-BirA*-HA-IRES-Neo vector.

The V5-BirA*-H2A.B was created by inserting BirA* between V5 and H2A.B within the pEF1a-V5-H2A.B-IRES-P construct and then cloning V5-BirA*-H2A.B into pLVX-EF1α-IRES-Neo vector using EcoRI and BamH1 restriction sites, generating the pLVX-EF1a-V5-BirA*-H2A.B-IRES-Neo vector. The lentiviral transduction was carried out to create HL cell lines stably overexpressing the above constructs as described for H2A.B knockdown cell line establishment with 20αg of V5-BirA*-H2A.B or H2A.B-BirA*-HA vectors in place of a SMARTvector. FACS sorting and doxycycline (dox) induction was not required.

### Cell proliferation assay

The cell proliferation assay was performed in sterile 96-well round bottom microwell plates using CyQUANT direct cell proliferation assay (Thermo Fisher Scientific). The assay was performed on ~ 5,000 of L1236 or L428 cells in 100 μl of growth medium per well, in quintuplicates, and analysed on a FLUOstar OPTIMA plate reader (BMG LABTECH).

### KCl-based subcellular fractionation

The fractionation was performed using 1 x10^7^ cells as previously described (Soboleva et al., 2017).

### Western blot

Protein samples were transferred onto a immobilon-PSQ 0.2 μM pore size PVDF membrane (Merk Millipore). The PVDF membrane was blocked by 3% BSA in 0.05% PBST (0.05% Tween-20 in 1 x PBS) for 1h at room temperature. The membrane was incubated with the primary antibody diluted in 1% BSA/PBST overnight at 4°C. The secondary antibody, conjugated with horseradish peroxidase, was incubated with the membrane for 1 h at room temperature.

### Immunofluorescence and microscopy

Cells (2×10^6^) were treated in incubated in ice-cold hypotonic solution (0.1 M sucrose, Tris-HCl pH 8.1, and 1 x EDTA-free protease inhibitor cocktail) on ice for 6 min. Nuclei were then fixed on poly-L-lysine coated glass in 50-60 μL of fixation/permeabilization solution (2% paraformaldehyde, 0.1% Triton X-100, pH 9.2) in a humidified chamber for 2 h at room temperature. After fixation, the slides were air-dried and used for immunofluorescence staining. The slides were blocked by 3% BSA in DPBS for 1 h, at room temperature. Primary and secondary antibodies were diluted typically to 1/100-1/200 (v/v) in Antibody Dilution buffer (1% BSA in 0.1% PBST). The primary antibody probing was performed at 4°C, overnight followed by three washes in DPBS, followed by secondary antibody probing for 45 min at 37°C. The nuclei were counterstained with 0.4 μg/ml DAPI in water and samples were mounted with VECTASHIELD medium (Vector laboratories) and sealed with cover slips. The slides were imaged using the Leica SP5 confocal microscope (Leica Camera). For live cell imaging, the cells in 6-well plates or flasks were imaged directly using the Olympus IX71 inverted fluorescence microscope (Olympus Corporation).

### Immunoprecipitation (IP) and chromatin Immunoprecipitation (ChIP)

20 μl of protein A and protein G Dynabeads each (Thermo Fisher Scientific) slurry were washed twice in 900 μl beads wash buffer (0.02% Tween-20 in DPBS) and incubated with antibody (typically 5 μg) for 1 h on a wheel at RT in 200 μl beads wash buffer. To remove the unbound antibody, the beads were washed twice in 900 μl beads wash buffer.

For IP, whole cell lysate was prepared from 10 x10^6^ cells by lysing them in 1 ml iCLIP lysis buffer (50 mM Tris-HCl pH 7.4, 100 mM NaCl, 1% NP-40, 0.1% SDS, 0.5% sodium deoxycholate, 1 mM AEBSF and 1 x EDTA-free protease inhibitor cocktail). The lysate was homogenized by passing 5 - 10 times through a various gauge needles, and sonication using the Bioruptor® (Diagenode) for 15 cycles of 30 s ON / 30 s OFF at high setting. After sonication, the lysate was centrifuged for 30 min at 16,000 x g at 4°C. The supernatant was subjected to IP by incubating it with antibodies bound to Protein A/G Dynabeads for 4h to overnight at 4°C. The beads were washed 3 times in 1 ml iCLIP lysis buffer, twice in 1ml iCLIP High Salt buffer (iCLIP lysis buffer supplemented with 1 M NaCl), and twice in 1 ml iCLIP Wash buffer (20 mM Tris-HCl, 10 mM MgCl2, 0.2% Tween-20, pH 7.4). IP samples were eluted from the beads by incubating in LDS loading buffer (Life Technologies) at 80°C for 10 min and analysed by Western blot. ChIP assays and the preparation of ChIP-Seq libraries were performed as previously described by us (Soboleva et al., 2012; Soboleva et al., 2017).

### Cleavage Under Targets and Release Using Nuclease (CUT&RUN) followed by sequencing

The CUT&RUN procedure was performed as previously described (Skene et al., 2018), with minor modifications. 4μg of H2A.B antibody was diluted into 200 μl icecold Dig-wash buffer (20 mM HEPES -NaOH, 150 mM NaCl, 0.5 mM spermidine, 1 x protease inhibitor cocktail, and 0.1% digitonin, pH 7.5) supplemented with 2 mM EDTA and incubated with 10 μl of ConA beads (Bangs laboratories) slurry and 0.25 x10^6^ cells o/n on a wheel at 4°C. For each experiment, 1 μg of H3K36me3 antibody (positive control) and 4μg of rabbit IgG (negative control) antibody were used in parallel.

Following antibody binding, the cells were washed twice in 1 ml Dig-wash buffer and resuspended in 200 μl pA/MNase solution, gift from Henikoff laboratory, (~ 350 ng/ml in Dig-wash buffer), followed by incubation for 1 h on a wheel at 4°C. and washed twice in 1 ml Dig-wash buffer. The cell pellets were resuspended in 100 μl Dig-wash buffer, and placed into the metal block on ice to cool down to 0°C. 2 μl of 100 mM CaCl2 solution was mixed into each sample to activate the MNase digestion. The digestion was carried out by incubating for 30 min at 0°C. After incubation, each sample was added with 100 μl of 2 x Stop solution (340 mM NaCl, 20 mM EDTA, 4 mM EGTA, 0.02% Digitonin, 150 μg/ml GlycoBlue™, 25 μg/ml RNase, and 25 pg/ml Drosophila spike-in DNA). For DNA fragment release, the sample was incubated in the thermal mixer for 10 min at 450 RPM, 37°C. The supernatant was subjected to DNA purification by mixing with 2 μl of 10% SDS and 2.5 μl proteinase K (20 mg/ml), followed by incubation in the thermal mixer for 10 min at 700 RPM at 70°C and phenolchloroform DNA extraction and precipitation with 1 μl GlycoBlue™ (15 mg/ml) and 750 μl of absolute ethanol. The DNA was analysed by Agilent 2100 bioanalyzer prior to library preparation. The sequencing libraries were prepared using NEBNext Ultra II DNA library prep kit for Illumina following the kit manual with modification. During the PCR amplification step, 12 PCR cycles were performed, with a combined annealing and extension step for 10 s, to minimize the large DNA fragments. The pooled library from all samples were sequenced by the Illumina NextSeq 500 sequencer, using 75 cycles of paired end reads.

### Assay for Transposase-Accessible Chromatin using sequencing (ATAC-seq)

ATAC-seq was performed as previously described (Corces et al., 2017) using 50,000 cells. Tn5 transposase from the Illumina Nextera DNA library kit was used. All ATAC-seq libraries were quality-checked on an Agilent high sensitivity DNA chip using the Agilent 2100 bioanalyzer to detect a nucleosome ladder pattern. Sequencing was performed a NovaSeq 6000 (Illumina) sequencer in a 2 x 50 bp paired-end configuration.

### Proximity-dependent biotin identification (BioID)

The BioID H2A.B constructs were first validated by fluorescence microscopy of the biotin-treated cells, which were stained by streptavidin-Cy5. The BioID procedure was performed as previously described (Roux et al., 2013), with minor modification. The interacting proteins were biotin-labelled by growing cells (~ 0.5 x10^6^ cells/ml) in RPMI-1640 growth medium supplemented with 50 μM biotin for 24 h. The whole cell lysate was prepared by lysing cells in iCLIP Lysis buffer (50 mM Tris-HCl, 100 mM NaCl, 1% NP-40, 0.1% SDS, 0.5% sodium deoxycholate, 1 mM AEBSF and 1 x EDTA-free protease inhibitor cocktail, pH 7.4) aided by homogenization by passing the cells through various gouge needles and sonication using the Bioruptor® sonication system (Diagenode) for 15 cycles of 30 s ON / 30 s OFF at high setting. 100 μL Dynabeads™ MyOne^TM^ Streptavidin C1 beads were used per 1 ml cell lysate, adjusted to 2 mg/ml, for pull-down at 4 °C on a wheel overnight. After binding, the beads were washed twice in 1 ml Wash buffer I (2% SDS), once in Wash buffer II (50 mM HEPES-NaOH, 500 mM NaCl, 1 mM EDTA 1% Triton X-100, 0.1% sodium deoxycholate, pH 7.5), once in 1 ml Wash buffer III (10 mM Tris-HCL, 0.1% sodium deoxycholate, 0.5% NP-40, 1 mM EDTA, 250 mM LiCl, pH 7.4), and twice in 1 ml 50 mM Tris-HCl pH 7.4. The bound proteins were eluted with 100 μl of LDS sample buffer (Thermo Fisher) at 98°C.

### RNA-seq and differential expression analysis

After trimming the adaptor sequences using Trimmomatic, the RNA-seq samples in three replicates from the wild type and H2A.B mutant in two cell lines, L1236 and L428, were mapped to the *Homo sapiens* (hg38) genome assembly using HISAT2 (Kim et al., 2015) Gene annotation was obtained from the UCSC hg38 gene annotation in iGenomes. The sequencing reads were assigned to genes by featureCounts in Rsubread package in Bioconductor (Liao et al., 2019). Differentially expressed mRNAs between mutants versus wild type were identified, and FDR (Benjamini-Hochberg) was estimated, using DEseq2 (Love et al., 2014). The genes with FDR < 5% were considered to be significantly differentially expressed. The genes with an average gene expression log2 transcript per million (TPM) >3 were defined as expressed genes, which were used for the downstream analysis.

### Differential alternative splicing analysis

FeatureCounts was used to assign the RNA-seq sequencing reads to exons using a custom-made data table, where unique exons per transcript per gene were described. DSEXseq R package was used to identify differential exon usage between wild type and H2A.B mutant (Anders et al., 2012). The exons with FDR ≤ 10% were considered to be significantly differentially spliced exons.

### Exon Intron Split Analysis (EISA)

EISA was carried out as described by (Gaidatzis et al., 2015) with a modified custom script. In brief, only non-overlapping genes were included for the analysis. To ensure sufficient intronic and exonic counts, we applied non-specific filtering requiring the genes to have normalised counts to sequencing depth greater than a threshold for both intron and exon (mean(log2(nomalised-counts + 8)) > 5). A modified custom script was incorporated into edgeR package (Robinson et al., 2010) to identify genes (FDR < 0.05) with a significant difference in the level of exon, intron, or exon minus intron between H2A.B knock-down versus control.

### CUT&RUN bioinformatic analysis

CUT&RUN of H2A.B wild-type and IgG samples were performed in three replicates with *Drosophila melanogaster* cell chromatin as spike-in. The CUT&RUN samples were mapped to the *Homo sapiens* (hg38) genome assembly using Bowtie2 with the default parameters, after the adaptor trimming by Trimmomatic. The high quality and uniquely mapped reads with a mapping quality MAPQ > 20 were used for further analysis. The spike-in reads were obtained by mapping the reads to the *Drosophila melanogaster* (dm6) genome assembly. Generated coverage tracks for CUT&RUN samples, which were normalized to spike-ins. Peak calling of CUT&RUN reads was performed against IgG reads by Genrich peak caller (https://github.com/jsh58/Genrich), which took the replicate consistency into account. The peak annotation was performed by HOMER package (Heinz et al., 2010). Peak overlap analysis was performed by “mergePeaks” function in HOMER package with the default parameters. The pathway enrichment analysis of 404 genes genes containing H2A.B peaks within 2Kb of the TSS and commonly down-regulated in L1236 and L428 HL cell lines in response to H2A.B knockdown were performed by STRING (Szklarczyk et al., 2019).

### ATAC-seq analysis

The ATAC-seq samples in two biological replicates in two cell lines were mapped to the *Homo sapiens* (hg38) genome assembly using Bowtie2 with default parameters after adaptor trimming by Trimmomatic. The high quality and uniquely mapped reads (MAPQ > 20) and the reads filtered for PCR-duplicates by Picard were used for further analysis. We performed peak calling of ATAC-seq accessible regions by Genrich peak caller with parameter setting (Genrich -t -o -f -r -j -y -d 100 -q 0.05 -e chrM -v). The peaks were annotated using the HOMER package. ATAC-seq differential accessibility analysis between wild-type and H2A.B mutant on the CUT&RUN peaks was performed with the DEseq2 R package.

## DATA AVAILABILTY

GEO accession GSE158239.

## AUTHOR CONTRIBUTION

X.J. designed and performed the experiments; J.W. performed the bioinformatic analysis; E.P. analysed HL primary samples; W.Y. produced and characterised H2A.B antibody; G.S. characterised HL cell lines and sequenced the H2A.B genes in HL cell lines; A.B. performed histological staining of HL primary sample; J.E.D. analysed HL primary samples and edited the manuscript; D.J.T. and T. A.S. designed and supervised the study and wrote the manuscript.

## ACKNOWLEDGEMENTS

We thank Steve Henikoff for a gift of pA/MNase; Sebastian Kurschield for the initial bioinformatic analysis, Ross Hannan and Kate Hannan for helpful discussions and provision of anti-RNA Pol-I antibodies and primers for qPCR. We especially thank Radjiv Khanna who provided us with the HL cell lines.

